# Characterising pre-clinical sub-phenotypic models of Acute Respiratory Distress Syndrome: an experimental ovine study

**DOI:** 10.1101/2020.12.02.408682

**Authors:** Jonathan E Millar, Karin Wildi, Nicole Bartnikowski, Mahe Bouquet, Kieran Hyslop, Margaret R Passmore, Katrina K Ki, Louise E See Hoe, Nchafatso G Obonyo, Lucile Neyton, Sanne Pedersen, Sacha Rozencwajg, J Kenneth Baillie, Gianluigi Li Bassi, Jacky Y Suen, Daniel F McAuley, John F Fraser

## Abstract

The Acute Respiratory Distress Syndrome (ARDS) describes a heterogenous population of patients with acute severe respiratory failure. However, contemporary advances have begun to identify distinct sub-phenotypes that exist within its broader envelope. These sub-phenotypes have varied outcomes and respond differently to several previously studied interventions. A more precise understanding of their pathobiology and an ability to prospectively identify them, may allow for the development of precision therapies in ARDS. Historically, animal models have played a key role in translational research, although few studies have so far assessed either the ability of animal models to replicate these sub-phenotypes or investigated the presence of sub-phenotypes within animal models. Here, in three ovine models of ARDS, using combinations of oleic acid and intravenous, or intratracheal lipopolysaccharide, we demonstrate the presence of sub-phenotypes which qualitatively resemble those found in clinical cohorts. Principal Components Analysis and partitional clustering reveal two clusters, differentiated by markers of shock, inflammation, and lung injury. This study provides the first preliminary evidence of ARDS phenotypes in pre-clinical models and develops a methodology for investigating this phenomenon in future studies.

## Introduction

It is increasingly understood that the Acute Respiratory Distress Syndrome (ARDS) describes a clinically and immunologically heterogenous population (1). Heterogeneity among patients with ARDS has been proffered as an explanation for consistently negative trials of pharmacological treatments. Contemporary advances in phenotyping, using unsupervised machine learning techniques, have identified novel sub-phenotypes in clinical trial cohorts (2). These phenotypes have discrepant outcomes, and importantly, appear to respond differently to several interventions (3). An ability to prospectively identify sub-phenotype membership in patients with ARDS opens the possibility of delivering personalized treatments.

Historically, animal models of ARDS have played an important role in biological discovery and in therapeutic translation (4). Numerous models of ARDS have been developed in both large and small animals. However, an animal model that fully recapitulates the clinical pathobiology of ARDS is not available. This has contributed to the gap between results generated from pre-clinical models and those obtained in subsequent clinical trials. As our knowledge of clinical sub-phenotypes grows, a new question arises for those modelling ARDS in animals; how well does an animal model reflect the pathobiology of a specific clinical sub-phenotype? To answer this question several preliminary facts need to be elucidated. Do existing pre-clinical models of ARDS more closely resemble one phenotype or another? And, do animals with experimental ARDS exhibit phenotypes given a common method of injury?

Thus, we sought to develop an approach to these problems by testing three models of ARDS in sheep. Using a combination of dimensionality reduction and partitional clustering, we investigated the presence of sub-phenotypes, arising dependent or independent of the means of injury. Previously, others have pursued a related approach to identify sub-phenotypes in a murine model of sepsis (5). Similarly, we aimed to provide preliminary evidence of distinct sub-phenotypes arising in pre-clinical models of ARDS, and to propose a methodology for investigating these phenomena in animal models.

## Materials and methods

### Study design

Ethical approval for this study was obtained from University Animal Ethics Committees (QUT1600001108, UQPCH/483/17). The study was conducted in accordance with the Australian Code of Practice for the Care and Use of Animals for Scientific Purposes (6), and reported in compliance with the ARRIVE guidelines (7). Detailed methods are provided in an online supplement. A diagrammatic summary of the study is presented in **Figure 1**.

**Figure 1.**
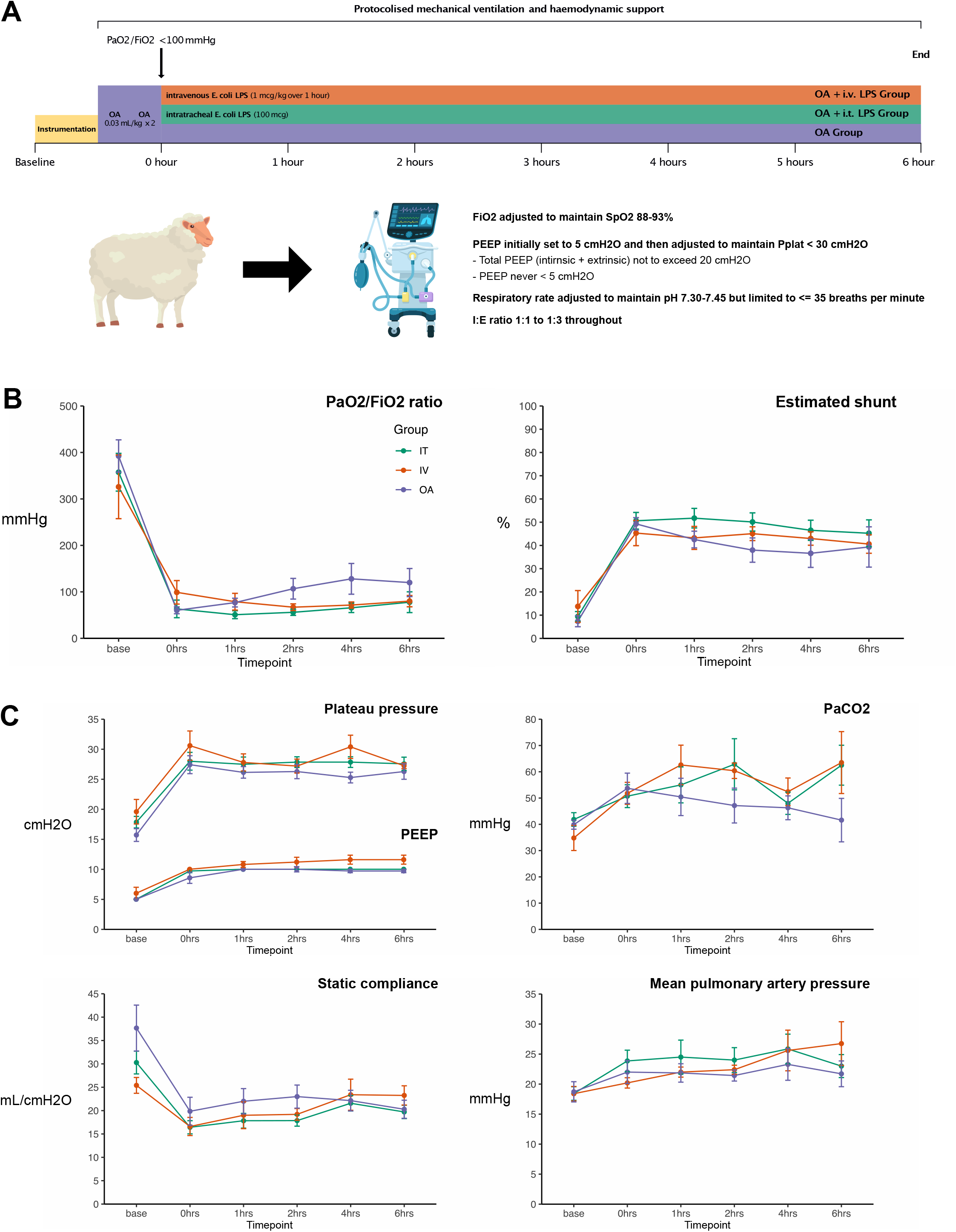
Study overview, measures of gas exchange, and respiratory mechanics. A. Schematic overview of study design. B. Measures of gas exchange. C. Measures of respiratory mechanics. Data are presented as mean and 95% confidence intervals.

### Animal model

Nineteen healthy Border Leicester Cross ewes, aged between 1-3 years and weighing 52 kg (47-54), were randomly assigned to one of three groups; injury by intravenous infusion of oleic acid (OA, n=7), by OA and intratracheal *E. coli* lipopolysaccharide (IT, n=7), or by OA and intravenous *E. coli* lipopolysaccharide (IV, n=5).

Briefly, animals were anesthetized with ketamine, midazolam, and fentanyl. Continuous neuromuscular blockade was maintained by infusion of vecuronium. After induction, animals were tracheostomized and ventilated using a low tidal volume strategy. After instrumentation, acute lung injury was induced by infusion of OA (0.06 mL/kg; O1008, Sigma-Aldrich, Castle Hill, NSW, Australia), with or without, intratracheal *E. coli* LPS (100 μg; O55:B5, Sigma-Aldrich, Castle Hill, NSW, Australia) or intravenous *E.coli* LPS (1 μg/kg infused over one hour; O55:B5, Sigma-Aldrich, Castle Hill, NSW, Australia). Once a PaO_2_/FiO_2_ ratio <100 mmHg (PEEP ≥5 cmH_2_O) was achieved (0 hours), animals received protocolized intensive care for the duration of the study. At six hours, animals were euthanized, and lung tissue was recovered for examination.

### Blood and cytokine analysis

Blood samples were analyzed by and independent veterinary laboratory (IDEXX Laboratories, Brisbane, Australia) to clinical standards. Routine biochemical, hematological, and coagulation tests were performed. The concentration of interleukin-6 (IL-6), IL-8, IL-1β, and IL-10 were measured in blood and in bronchoalveolar lavage (BAL) fluid. Our development of ovine specific ELISA assays has been described before (8). Detailed methods are provided in an online supplement.

### Statistical analysis

Data are expressed as median (IQR). Analysis was undertaken in R 4.0.3 (R Core Team, Vienna, Austria, 2020). The dataset for this study, along with reproducible code, is available at https://github.com/JonathanEMillar/Characterising-pre-clinical-sub-phenotypic-models-of-ARDS. Longitudinal data were analyzed by fitting a linear mixed model, using the R package *lme4*. Nonlongitudinal data were compared with one-way ANOVA. Where a significant interaction was observed, *post-hoc* comparisons were made using Tukey’s test, using the R package *rstatix*. Correction for multiple comparisons was made using the Benjamini-Hochberg method. Frequency data were compared using the chi-squared test. Co-linearity was assessed by calculating the Spearman correlation coefficient for each pair of variables, using the R package *corrplot*. Principal component analysis (PCA) was performed to reduce the dataset to a smaller number of principal components (PCs), using the R package *FactoMineR* (9). Beforehand, missing data were imputed using a random forest approach with predictive mean matching (using the R package *missRanger*) and the dataset was z-score normalized by subtracting the variable mean and dividing by the variable standard deviation (10). After examination of the scree plot, principal components sufficient to explain >75% of the total variance were retained. Partitioning Around Medoids (PAM) clustering was performed, after PCA, on the imputed dataset, using the R package *cluster*. A Euclidean distance measure was employed. The optimal number of clusters to specify was derived from a ‘majority’ assessment of 26 measures, using the *NbClust* package. In event of a tie a parsimonious solution was preferred. To assess the stability of clusters, we employed a non-parametric bootstrap-based strategy using the R package *fpc*. This generated 1000 new datasets by randomly drawing samples from the initial dataset with replacement and applying the same clustering technique to each. Clustering results were then compared for each cluster identified in the primary analysis and the most similar cluster identified for each random re-sampling. A mean value for the Jaccard coefficient, for the sum of the comparisons, was generated for each cluster. Z-scores for each variable, by cluster, were descriptively compared with clusters derived from a previously published latent class analysis of the ARMA study (2), obtained using a digital ruler. Statistical significance was assumed if *P* < 0.05.

## Results

Baseline characteristics at injury (0 hours) are summarized in **Table 1** and in **Supplementary Table E1**. All animals completed the study protocol and were euthanized at six hours.

**Table 1.** Physiological characteristics at zero hours (injury). Data are presented as median (IQR).

|  | **Overall**<br>(n=19) | **OA**<br>(n=7) | **IT**<br>(n=7) | **IV**<br>(n=5) |
| --- | --- | --- | --- | --- |
| **Weight** (kg) | 52 (47-54) | 55 (53-57) | 47 (46-51) | 52 (46-52) |
| **PEEP** (cmH<sub>2</sub>O) | 10 (10-10) | 10 (7.5-10) | 10 (10–10) | 10 (10-10) |
| **Plateau pressure** (cmH<sub>2</sub>O) | 27 (26-30) | 26 (26-27) | 27 (26-29) | 29 (28-34) |
| **Static compliance** (mL/cmH<sub>2</sub>O) | 16 (14-22) | 21 (16-25) | 16 (14-18) | 16 (14-16) |
| **PaO<sub>2</sub>/FiO<sub>2</sub>** (mmHg) | 52 (47-90) | 52 (49-69) | 47 (46-51) | 100 (55-133) |
| **Effective shunt** (%) | 49 (43-54) | 49 (44-54) | 50 (48-54) | 45 (37-52) |
| **PaCO<sub>2</sub>** (mmHg) | 49 (46-57) | 47 (46-53) | 52 (46-57) | 49 (48-53) |
| **pH** | 7.31 (7.28-7.35) | 7.31 (7.25-7.32) | 7.33 (7.3-7.38) | 7.32 (7.31-7.37) |
| **Bicarbonate** (mmol/L) | 23.8 (22.3-24.9) | 22.4 (21.9-23.3) | 24.3 (23.9-24.9) | 23.4 (22.6-25.4) |
| **Base excess** (mmol/L) | -0.6 (-2.5-1.1) | -2.5 (-3- -2.4) | 1 (0-1.8) | 0.8 (-1.4-2.6) |
| **Heart rate** (bpm) | 112 (102-134) | 121 (96-134) | 111 (100-132) | 112 (106-117) |
| **Mean arterial pressure** (mmHg) | 96 (85-109) | 104 (96-111) | 80 (76-92) | 108 (99-115) |
| **Mean pulmonary arterial pressure** (mmHg) | 21 (19-25) | 22 (19-26) | 23 (20-27) | 20 (19-21) |
| **Central venous pressure** (mmHg) | 12 (9-13) | 13 (12-15) | 11 (9-13) | 11 (3-19) |

### Oleic acid, with or without LPS, induces acute severe respiratory failure. However, LPS prolongs or prevents improvements in oxygenation in the first 6 hours

The infusion of OA produced severe lung injury (PaO_2_/FiO_2_ <100 mmHg). However, oxygenation began to recover within one hour (**Figure 1**). LPS consistently maintained PaO_2_/FiO_2_ within the range of severe ARDS. Differences in oxygenation between groups were not associated with differences in respiratory system mechanics (**Figure 1**).

### The addition of LPS increases the plasma concentration of interleukin-6

Both intravenous and intratracheal LPS increased the plasma concentration of IL-6 (group:time, p <0.001) when compared to OA alone (**Supplementary Figure E1 and Supplementary Tables E3 and E4**). The use of intravenous LPS was associated with non-sustained increases in the plasma concentration of IL-8 (group:time, p <0.001). Similarly, the use of intravenous LPS resulted in increases in plasma IL-10 (group:time, p <0.001). All three injury methods were associated with low plasma levels of IL-1β. Cytokine concentrations in bronchoalveolar lavage (BAL) fluid were less distinct between groups (**Supplementary Figure E1 and Supplementary Tables E3 and E4**). Animals injured with intravenous LPS, as compared to OA alone, had higher white cell and neutrophil counts at 6 hours (p = 0.049 and 0.034 respectively).

### In the first six hours, intravenous LPS induces more severe shock in comparison to intratracheal LPS or OA alone

All pulmonary injury methods produced shock requiring vasopressor support (**Supplementary Figure E2**). While reductions in MAP from baseline to 6 hours were greater in animals receiving LPS, these differences were not statistically significant. Similarly, there were no significant differences in heart rate or CVP, between groups (**Supplementary Figure E2**). Cumulative fluid balance was greatest in OA animals, and significantly different when compared to sheep given LPS (**Supplementary Figure E2**). Cumulative urine output did not differ between groups (p = 0.109).

Hematological and biochemical parameters, at 6 hours, are presented in **Supplementary Table E2**. Following correction for multiple comparisons, there were no significant differences between groups in indices of renal function.

### Differences in variables describing the severity of shock and of lung injury explain the majority of variance between animals

There was a high degree of correlation between variables recorded in our study at 6 hours (**Supplementary Figure E3**). In an effort to reduce the dimensionality of the data and eliminate colinearity we performed PCA. There were some missing data in our study (**Supplementary Table E5**) and prior to analysis these data were imputed using a random forest method (**Supplementary Table E5**). In total, 8 principal components (PCs) explained > 75% of the variance between animals, with the first 4 PCs explaining > 50% (**Figure 2**). Markers of shock severity contributed heavily to PCs 1 and 2, while markers of lung injury defined PCs 4 and 5 (**Figure 2 and Supplementary Figure E4**).

**Figure 2.**
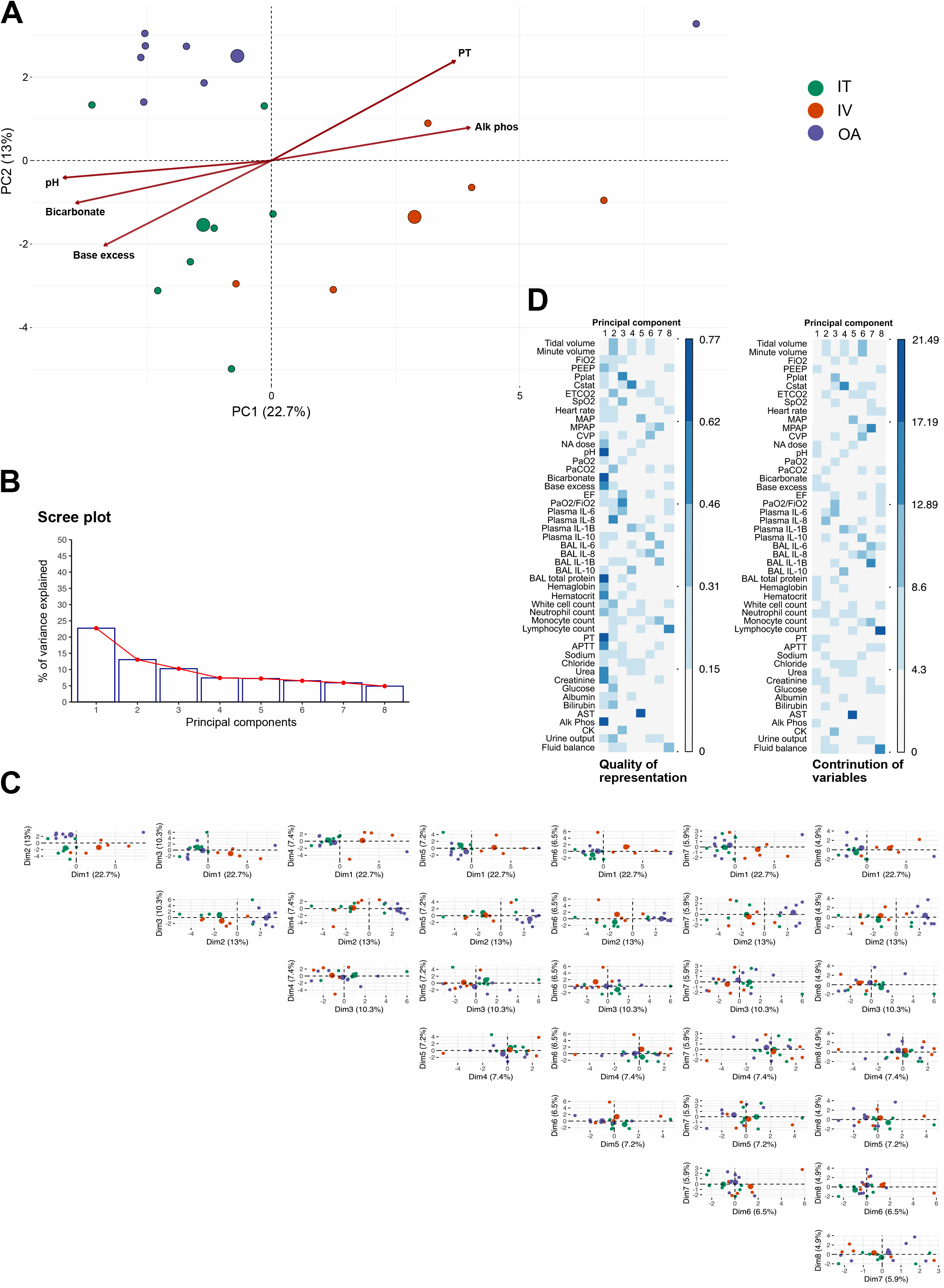
Principal components analysis (PCA). A. Biplot of principal components (PCs) 1 and 2. The top 5 variables in PC 1 are shown. Large dots represent the group mean. B. Scree plot of first 8 PCs. C. Pairs plot of PCA projections for first 8 PCs. D. Contribution and quality of representation of variables to PCs. The quality of representation (cos2) sums to 1 for each variable across all PCs. The contribution of variables to variance in a PC are expressed as percentages.

### Two distinct phenotypes are evident and are associated with the method of injury

An *a priori* assessment of the data suggested that two clusters was the optimal partition of our data (**Supplementary Table E6**). The distribution of the clusters projected onto PCs 1 and 2 of the PCA is shown in **Figure 3**. Animals in cluster B (n = 4) tended to have a lower MAP, lower urine output, were more acidotic, coagulopathic, and had higher plasma levels of IL-6, IL-8, and IL-10 (**Figure 2 and Figure 4**). Proportionally, more animals injured with intravenous LPS were found in Cluster B (p = 0.036). All animals injured with IT LPS belonged in Cluster A. The clusterwise stability was good (mean Jaccard similarities, 0.85 Cluster A, 0.71 Cluster B). Next, we aimed to compare the characteristics of the clusters to those identified in an ARDS clinical trial cohort. We extracted z-scores for overlapping variables from a latent class analysis of the ARMA study (2) and plotted the hyper-inflammatory cohort against Cluster B and the hypo-inflammatory cohort against Cluster A (**Figure 4**). There was general concordance between both pairs, with the exception of minute volume, bilirubin, sodium, and glucose levels.

**Figure 3.**
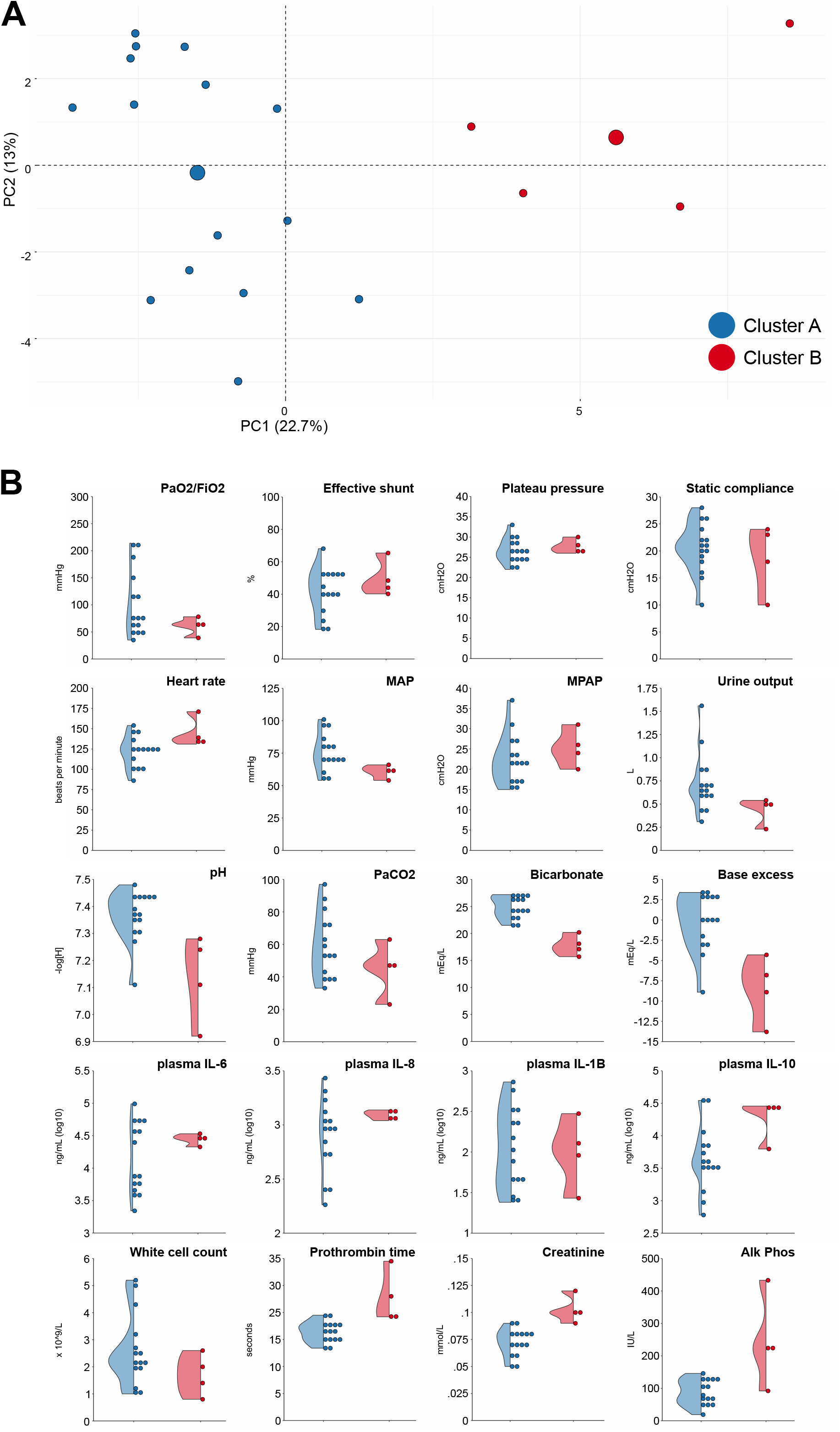
Partitional clustering and sub-phenotypes. A. Clustering results projected on PCs 1 and 2 of the PCA. B. Half-dot, half-violin plots of key variables stratified by cluster membership.

**Figure 4.**
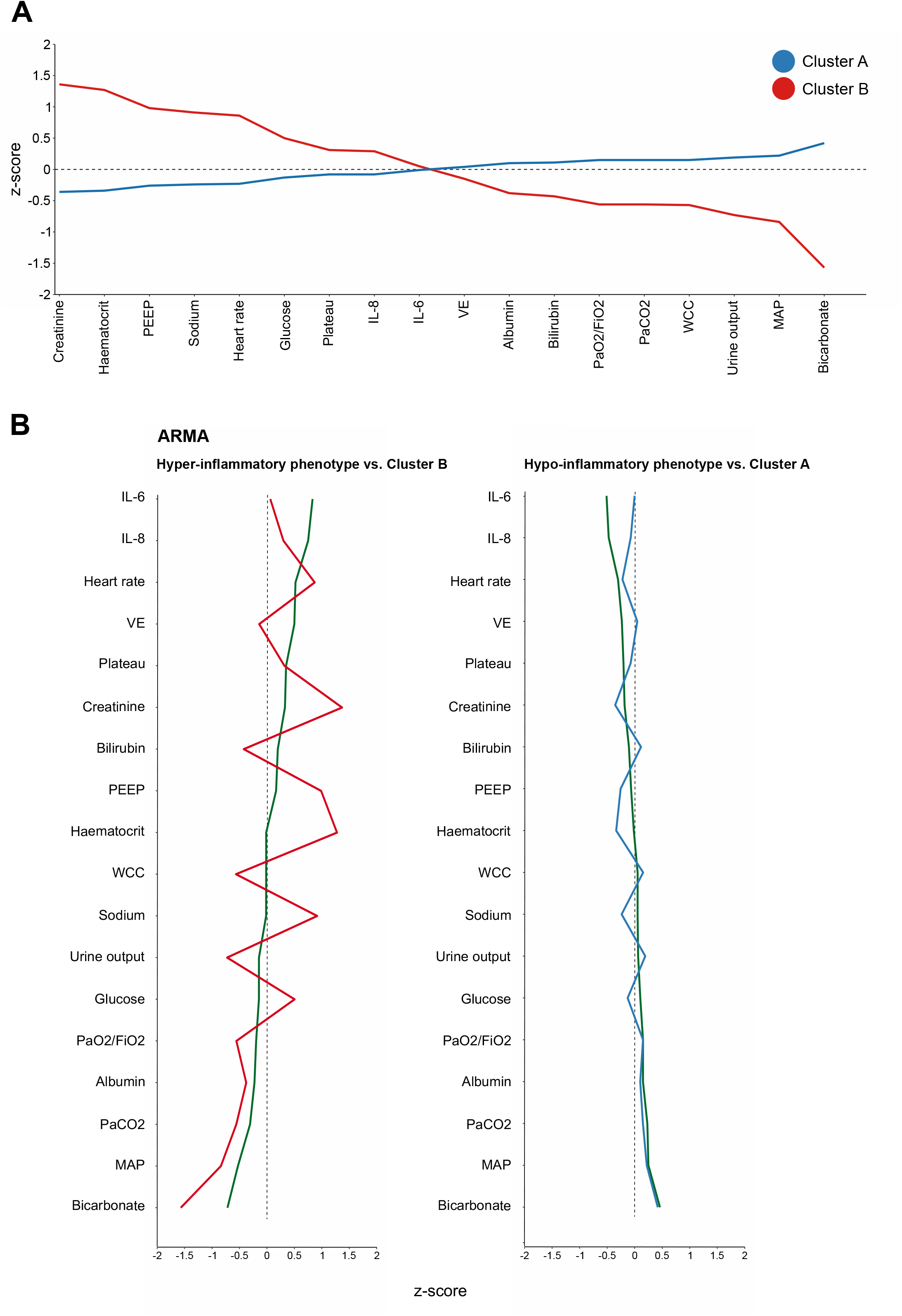
Pre-clinical clusters and clinical cohorts. A. Z-core plot of animal clusters for variables common with those in published clustering studies. VE = minute volume, WCC = white cell count, MAP = mean arterial pressure. B. Pre-clinical cluster z-score plots contrasted with clinical trial clustering sub-phenotypes. Green line = ARMA trial sub-phenotypes.

## Discussion

In a study combining three ovine models of ARDS, we provide preliminary evidence of subphenotypes occurring early in the course of illness. In doing so, we provide the first characterisation of OA and LPS ‘double-hit’ models of ARDS in large animals. In this study, OA was capable of inducing severe acute respiratory failure, however oxygenation began to improve rapidly unless animals were additionally insulted with LPS. All three injury models produced an elevation in plateau airway pressure and a reduction in compliance. Likewise, all exhibited evidence of shock, requiring vasopressor support. Plasma levels of IL-6 were higher in animals receiving intravenous LPS, however BAL concentrations remained unchanged. Decomposition of the dataset by PCA, revealed that the first 4 PCs explained 54% of the variance in the dataset, and may be characterised as representing the degree of shock/inflammation (PCs 1 and 2) and lung injury (PCs 3 and 4). Unsupervised clustering, using PAM, identified the presence of two distinct sub-phenotypes. These were primarily differentiated along PC1. Cluster membership was associated with method of injury with more animals belonging to the intravenous LPS group in cluster B. These data may have implications for translational research in ARDS. First, the presence of sub-phenotypes in animal models opens the possibility of developing phenotype-specific models. Second, unappreciated sub-phenotypic variation in existing or future pre-clinical studies may result in differential treatment effects where interventions are being studied.

OA infusion is a classical model of ARDS, first described by David Ashbaugh and colleagues, in 1971 (11). The intravenous infusion of OA results in rapid-onset lung injury, largely due to pulmonary endothelial damage and the formation of proteinaceous oedema in alveoli (12). This is evident in our data, where OA infusion generated PaO_2_/FiO_2_ ratios <100 mmHg, within 30 minutes. OA has also been associated with the upregulation of inflammatory cytokines and chemokines, such as; IL-6, IL-8, TNF-α, and matrix metalloproteinases (13, 14). However, OA is not implicated in the activation of several signalling pathways known to be important in clinical ARDS, such as NF-κB (15). In this study, animals receiving OA alone had consistently lower levels of pro-inflammatory cytokines measured in plasma. These differences may also be accounted for by the fact that the injury caused by OA is predominantly pulmonary, as 85% of the free-fatty acids from OA are retained in the lung (16).

These features may restrict the ability of OA to accurately replicate the full pathobiology of ARDS, particularly ARDS of a non-pulmonary aetiology. The addition of LPS may address some of these limitations. As a common constituent of Gram-negative bacterial cell walls, LPS participates in the pathology of causes of direct and indirect ARDS. The mechanisms of LPS-mediated injury include; lung epithelial injury (involving NF-κB induction), pulmonary endothelial damage, neutrophilic infiltration, and the activation of alveolar macrophages (17). The intravenous infusion of LPS is also associated with a systemic inflammatory response and the development of shock in pre-clinical models; as an alternative, by administering LPS via the intratracheal route, systemic effects may be limited, while inducing similar pathways within the lung (18). In this study, the addition of LPS was associated with prolonged severe hypoxaemia, in contrast to OA alone.

An expanding number of studies have identified two consistent sub-phenotypes of ARDS in clinical cohorts, which have been broadly characterised as hyper- and hypo-inflammatory (2, 3, 19–21). These sub-phenotypes have distinct outcomes and, in retrospective analysis, have been shown to respond differently to several interventions. Related sub-phenotypes have also been identified in other conditions causing critical illness (22–24). There are methodological challenges in assessing the similarity of these sub-phenotypes between studies, exacerbated by a lack of understanding of the mechanisms underpinning their development and differences in the methods used to cluster groups. However, to date, no study has sought to identify these groupings in animal models of ARDS.

Given the high degree of correlation between many of the measured variables in this study, a useful first step in identifying sub-groups is undertaking PCA. This technique uses orthogonal transformations to create a smaller number of linearly uncorrelated variables, while minimising information loss (25). The first PC in this study explains 23% of the variance in the dataset. It is comprised of several variables which may be associated with the severity of shock, such as; acid-base status, coagulation, and renal function. Alkaline phosphatase (Alk Phos) was also a prominent contributor. Alk Phos is expressed by granulocytes during sepsis and is involved in the detoxification of LPS (26), and levels have previously been associated with the severity of severe sepsis and intensive care unit length of stay (27). The third and fourth PCs, which together explain 18% of the variance, are predominantly influenced by PaO_2_/FiO_2_, plateau pressure, estimated shunt fraction (EF), and static compliance. EF is a non-invasive means of inferring shunt fraction from arterial blood gas data and has been found to be have superior predictive ability to PaO_2_/FiO_2_ ratio (28).

Having implemented three distinct models of experimental ARDS, partitional clustering was used to investigate the presence of sub-phenotypes within the combined study. Partitional clustering is a common method to partition objects, in this case animals, into an optimal number of related clusters (29). The aim is to maximise homogeneity within clusters while minimising it between them. The algorithm is agnostic to the number of clusters, therefore an assessment of the optimal number of clusters was made before clustering was applied. In this study, two clusters provided the optimal solution by a majority of measures. Mapping of individual animals on to the PCs derived by PCA allows us to visualise the differences between clusters in a two-dimensional space.

The sub-phenotypes identified in this study share some qualitative similarities with those described in ARDS clinical cohorts. Cluster B animals in this study exhibited higher levels of plasma IL-6, IL-8, creatinine, and serum sodium, with elevated prothrombin times and heart rates. On the other hand, they had lower serum albumin, leukocyte count, bicarbonate concentration, and PaO_2_/FiO_2_ ratio. This is a similar trend to that of the hyper-inflammatory sub-phenotype identified in clinical studies (2). Advances in methodology alongside more granular phenotyping, using -omics technologies, may be required to provide a better estimation of the validity of any comparison.

This study has some important limitations. First, large animals, like humans, exhibit variability in their response to injury. This effect of variability can be reduced with the inclusion of a greater number of experimental subjects. This study used a small number of animals, in part due to the exploratory nature of the design, which limit the conclusions which can be drawn. However, large animal experimentation, particularly those involving complex critical care interventions, are resource intensive and future studies are likely to face challenges in including substantially greater numbers. Second, although differences were observed between groups and sub-phenotypes, the duration of the study was short. A longer period of sampling may have captured evolving differences among measured parameters. This may especially true of indices in which there is likely to be lag after injury, such as renal dysfunction. Third, the choice of clustering technique is flexible, given the variety available to investigators. PAM has several specific disadvantages, not least of which is its vulnerability to the influence of outliers. Future studies may seek to validate clustering solutions by adopting more than one technique (30).

In conclusion, we identified preliminary evidence of sub-phenotypes occurring in animal models of ARDS. These phenotypes are characterised by differences in the severity of shock, systemic inflammation, and lung injury. Sub-phenotypes bear a qualitative similarity to those identified in clinical cohorts. The method of injury chosen in animal models may tend toward one or the other. Further studies are required to confirm these findings and to develop our understanding of the biological underpinning of sub-phenotypes in ARDS.

## Authors contributions

J.E.M - study conception, model development, study design and conduct, animal surgery, data
collection and analyses, manuscript preparation.

K.W. - model development, study design and conduct, animal surgery, data collection and analyses, manuscript preparation.

N.B., M.R.P., J.Y.S. – model development, study design and conduct, data collection and analyses, manuscript review.

N.G.O., S.P., S.R. – model development, study design and conduct, animal surgery, manuscript review.

M.B., K.H., M.Y., K.K.K. – sample analyses, data analyses, data collection, manuscript review.

J.K.B. –1 data analyses, manuscript review.

G.L.B. - study design, manuscript review.

D.F.M., J.F.F. - study conception, model development, study design, manuscript review.

## Online data supplement

This article has an online data supplement, which is accessible from this issue’s table of content online.

## Online Supplement

### A. Sheep preparation

Healthy, farm reared, female sheep (*Ovis aries*, Border Leicester Cross), aged 1-3 years, were used in this study. Animals were allowed a minimum of two weeks acclimatisation at the research facility, during which time they were housed socially in an outdoor enclosure. Prior to experimentation animals underwent veterinary examination and were screened for common ovine pathogens. The day before inclusion, sheep were housed in an indoor pen and given free access to water. Fasting from solid feed was enforced in the 12 hours prior to induction.

Prior to the induction of general anaesthesia, a central venous catheter (Arrow, Teleflex Medical Australia, Mascot, NSW, Australia) was inserted into the right external jugular vein (EJV), and baseline blood samples were obtained. An 8 Fr percutaneous introducer sheath (Arrow, Teleflex Medical Australia, Mascot, NSW, Australia) was also inserted in the right EJV. Animals were then pre-oxygenated for 3 minutes using a facemask. General anaesthesia was induced as described below. Animals were positioned supine. A surgical tracheostomy was performed, and a size 9.0 Portex tracheostomy tube (Smith’s Medical Australia, Sydney, NSW, Australia) was inserted. Correct positioning was confirmed using a flexible video bronchoscope (aScope™, Ambu, Ballerup, Denmark). Thereafter, the tracheostomy was suctioned hourly using an in-line suction catheter.

### B. Supportive care protocol

Animals were managed throughout the study by experienced critical care practitioners. Expert veterinary advice was available to investigators at all times.

#### Anaesthesia/analgesia

General anaesthesia was induced using intravenous midazolam (0.5 mg/Kg, Pfizer, Sydney, NSW, Australia) and ketamine (5 mg/kg, Troy Laboratories, Sydney, NSW, Australia). Animals were intubated (size 9.0-10.0 ID Portex endotracheal tube, Smith’s Medical Australia, Sydney, NSW, Australia) and mechanically ventilated. Maintenance of anaesthesia/analgesia was achieved by intravenous infusion of midazolam (0.5-0.8 mg/kg/hr, Pfizer, Sydney, NSW, Australia), ketamine (5-7.5 mg/kg/hr, Troy Laboratories, Sydney, NSW, Australia), and fentanyl (5-10 mcg/kg/hr, Hameln Pharmaceuticals, Hameln, Germany). Total intravenous anaesthesia was maintained until euthanasia.

Once maintenance anaesthesia was established, animals were paralysed with an intravenous bolus of vecuronium (50 mcg/kg, Pfizer, Sydney, NSW, Australia). Adequacy of neuromuscular blockade (NMB) was assessed using the train of four (TOF) response to peripheral nerve stimulation. NMB was maintained through the course of the experiment by an intravenous infusion of vecuronium (50 mcg/kg/hr, Pfizer, Sydney, NSW, Australia).

#### Mechanical ventilation

Animals were ventilated according to a protocolized lung-protective strategy. In a volume-controlled mode, the ventilator was set to achieve a tidal volume (V_t_) of 6 mL/kg actual body weight (ABW). One of, a Hamilton Galileo (Hamilton Medical, Bonaduz, Switzerland) or a Puritan Bennett 840 (Puritan Bennett, Medtronic, Dublin, Ireland) ventilator was used for the full course of each study. FiO_2_ was adjusted to achieve peripheral oxygen saturations (SpO_2_) in the range 88-93%, hyperoxia was avoided. PEEP was initially set to 5 cmH_2_O and adjusted to maintain a plateau pressure < 30 cmH_2_O. Total PEEP (extrinisic + intrinsic) could not exceed 20 cmH_2_O and could not be reduced < 5 cmH_2_O. Respiratory rate (RR) was adjusted to target a pH 7.30-7.45 but limited to ≤ 35 breaths per minute. An I:E ratio between 1:1 and 1:3 was used throughout.

#### Fluid and electrolyte management

A maintenance infusion of crystalloid (1-2 mL/kg/hr, compound sodium lactate, Baxter, Sydney, NSW, Australia) was provided for the duration of the study. Fluid therapy was indicated at other times to support haemodynamics, in this case 100 mL boluses of compound sodium lactate were titrated to effect.

Electrolyte levels were assessed 4-hourly. Serum potassium levels were maintained > 3.5 mmol/L by i.v. infusion of 10-20 mmol potassium chloride (AstraZeneca, Sydney, NSW, Australia). All large volume gastric losses were returned via the orogastric tube.

#### Monitoring and haemodynamic management

All animals were monitored according to the ‘*Recommendations for standards of monitoring during anaesthesia and recovery 2015: Association of Anaesthetists of Great Britain and Ireland*’ (1), including; pulse oximetry, 3-lead electrocardiogram, and continuous waveform capnography. In addition, the central venous catheter (CVC) was transduced as a measure of central venous pressure.

Invasive arterial blood pressure monitoring was established by cannulation of the left facial artery (20G Leadercath arterial, Vygon, Écouen, France). Using the 8 Fr percutaneous sheath introducer positioned in the right EJV, a pulmonary artery catheter (PAC) was inserted (7.5 Fr Swan-Ganz CCOmbo, Edwards Life Sciences, Irvine, CA, USA). Positioning was confirmed by obtaining a satisfactory waveform. Continuous cardiac output monitoring (Vigilance monitor, Edwards Lifesciences, Irvine, CA, USA) was instituted, with cardiac index estimated by; cardiac output/BSA [weight (kg)^0.67^ x 0.0842]. A 14 Fr urinary catheter (Gildana Healthcare, Oakleigh, VIC, Australia) was inserted and connected to an hourly urometer to allow for accurate quantification of urine output. A 16 Fr orogastric tube (ConvaTec, Glen Waverley, VIC, Australia) was inserted and left on free drainage.

A mean arterial pressure ≥ 65 mmHg was targeted. In the face of sustained hypotension, repeated 100 mL boluses of compound sodium lactate (Baxter, Sydney, NSW, Australia) were given until; CVP ≥ 8 mmHg and/or the animal was no longer fluid responsive (failure to increase stroke volume > 10% by PAC). Thereafter, noradrenaline (80 mcg/mL in 5% dextrose, Hospira, Lake Forrest, IL, USA) was commenced at 80 mcg/min and the dose titrated at 5minute intervals.

#### Euthanasia

At the end of each study, animals were euthanised by i.v. injection of phenobarbitone (142.5 mg/kg, Aspen Pharma, Dandenong, NSW, Australia). After death was confirmed (absence of cardiac electrical activity, blood pressure, and cardiac output monitoring), organs were retrieved surgically. Animal carcasses were stored and subsequently disposed of by incineration.

### C. Model of experimental ARDS

After instrumentation, animals were injured in the supine position.

#### Oleic acid preparation

A total dose of 0.06 ml/kg OA was used. Firstly, 0.03 mL/kg OA (O1008, Sigma-Aldrich, Castle Hill, NSW, Australia) was suspended in 20 mL arterial blood and 150 IU porcine heparin (Pfizer, Sydney, NSW, Australia). This mixture was administered via the distal port of the right EJV central venous catheter, followed by a flush of 50 mL 0.9% sodium chloride (Baxter, Sydney, NSW, Australia). The animal could recover and after 15 minutes this procedure was repeated. When 15 minutes elapsed from the second dose, arterial blood gas analysis, at a minimum PEEP of 5 cmH_2_O, was used to confirm a PaO_2_/FiO_2_ ratio <150 mmHg.

#### *E. coli* lipopolysaccharide preparation

Immediately after a PaO_2_/FiO_2_ ratio <150 mmHg was confirmed, *E. coli* LPS was administered via one of two routes to animals assigned to these groups.

For animals assigned to the i.t. LPS group, 50 mcg *E. coli* LPS (O55:B5, Sigma-Aldrich, Castle Hill, NSW, Australia), diluted in 10 mL 0.9% sodium chloride (Baxter, Sydney, NSW, Australia), was administered, via a designated video bronchoscope, to each main bronchus.

For animals assigned to the i.v. LPS group, 60 mcg *E. coli* LPS (O55:B5, Sigma-Aldrich, Castle Hill, NSW, Australia) diluted in 50 mL 0.9% sodium chloride (Baxter, Sydney, NSW, Australia), was administered infused via the CVC at a rate of 1 mcg/kg/hr for one hour.

### D. Clinical measurements

Hemodynamic and ventilatory data (including data derived from the PAC) were continuously monitored and automatically recorded at 5-minute intervals using a data monitoring system (Solar 8000, GE Healthcare, Waukesha, WI, USA) coupled with custom software. Urine and oro-gastric outputs were recorded on a pre-piloted observation chart on an hourly basis.

### E. Blood and bronchoalveolar lavage fluid analyses

#### Blood

Whole blood was sampled from the facial artery catheter. Arterial blood gas analysis was undertaken on at least an hourly basis (ABL800 FLEX, Radiometer, Copenhagen, Denmark). At baseline, zero hours (injury), 1 hour, 2 hours, 4 hours, and 6 hours, blood was sampled for routine laboratory haematological and biochemical testing. Testing was undertaken by an independent veterinary laboratory (IDEXX Laboratories, Brisbane, Australia) to clinical standards. Blood was also sampled for plasma cytokine measurements. The concentration of IL-6, IL-1β, IL-8 and IL-10 in plasma and bronchoalveolar lavage (BAL) fluid was quantified by in-house ELISAs. Positive internal controls were used to ensure that inter- and intra-plate variability was < 10% and confirm the precision and accuracy of all ELISA assays.

#### Bronchoalveolar lavage fluid

BAL was undertaken by an experienced bronchoscopist using a video bronchoscope (aScope™, Ambu, Ballerup, Denmark). At each examination the right and left middle and lower lobes were sampled. Each lobe was injected with 20 mL sterile 0.9% sodium chloride (Baxter, Sydney, NSW, Australia) and gentle suction was applied. The lavage fluid was collected in a sterile universal container. BAL fluid was centrifuged, and the supernatant collected for ELISA analysis. BAL total protein content was measured using a BCA (bicinchoninic acid) protein assay kit (Pierce BCA protein assay, ThermoFisher, VIC, Australia).

### F. Statistical analysis

The following R packages were used in the analysis, all versions as of 2020-11-04: tidyverse; psych; FactoMineR; factoextra; lme4; missRanger; rstatix; cluster; corrplot; RColorBrewer; cowplot; naniar; gghalves; NbClust; fpc; officer; flextable; gtsummary.

### G. Results supplement

**Supplementary Table E1.**
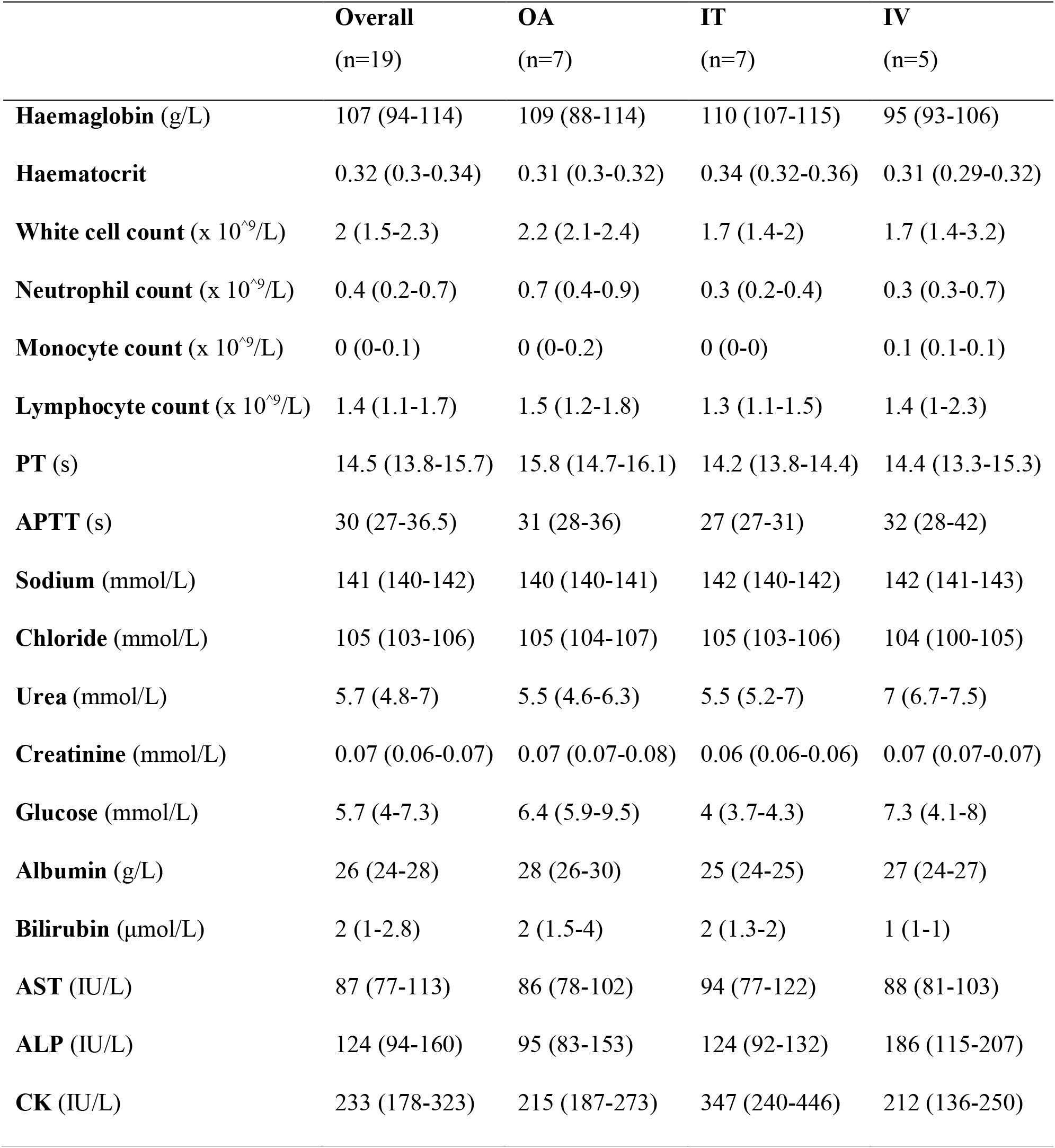
Hematological and biochemical characteristics at zero hours (injury). Data are presented as median (interquartile range).

**Supplementary Table E2.** Hematological and biochemical characteristics at 6 hours. Data are presented as median (interquartile range).

|  | **Overall**<br>(n=19) | **OA**<br>(n=7) | **IT**<br>(n=7) | **IV**<br>(n=5) |
| --- | --- | --- | --- | --- |
| **Haemaglobin** (g/L) | 100 (91-110) | 98 (91-109) | 94 (90-101) | 109 (105-114) |
| **Haematocrit** | 0.3 (0.27-0.35) | 0.28 (0.28-0.32) | 0.28 (0.26-0.31) | 0.34 (0.33-0.38) |
| **White cell count** (x 10<sup>9</sup>/L) | 2.2 (1.5-2.7) | 2.7 (2.2-3.8) | 2.1 (1.5-2.5) | 1.6 (1.1-2.2) |
| **Neutrophil count** (x 10<sup>9</sup>/L) | 1.3 (0.5-1.6) | 1.4 (1.4-2.7) | 1.2 (0.9-1.5) | 0.4 (0.3-0.5) |
| **Monocyte count** (x 10<sup>9</sup>/L) | 0.1 (0.1-0.2) | 0.1 (0.1-0.2) | 0.1 (0-0.2) | 0.1 (0.1-0.2) |
| **Lymphocyte count** (x 10<sup>9</sup>/L) | 0.8 (0.7-1.1) | 0.8 (0.8-1.1) | 0.8 (0.7-0.9) | 1.1 (0.7-1.5) |
| **PT** (s) | 17.6 (16.1-19.3) | 17.6 (17.2-18.5) | 15.2 (14.8-17) | 19.3 (19.3-21.2) |
| **APTT** (s) | 34.7 (29-37) | 33 (29-35) | 30 (29.5-38.5) | 37 (37-37) |
| **Sodium** (mmol/L) | 143 (140-144) | 142 (140-143) | 143 (140-143) | 144 (144-147) |
| **Chloride** (mmol/L) | 108 (105-109) | 107 (106-110) | 107 (105-108) | 109 (109-112) |
| **Urea** (mmol/L) | 5 (4.3-6.9) | 4.6 (4.3-5.2) | 5.2 (4.9-6.3) | 8 (3.9-8.2) |
| **Creatinine** (mmol/L) | 0.08 (0.07-0.09) | 0.08 (0.07-0.08) | 0.08 (0.06-0.08) | 0.09 (0.07-0.1) |
| **Glucose** (mmol/L) | 1.8 (1.4-2.8) | 2.7 (2.1-3.5) | 1.8 (1.3-2.2) | 1.5 (1.1-2.5) |
| **Albumin** (g/L) | 21 (19-24) | 25 (22-26) | 20 (20-21) | 19 (16-22) |
| **Bilirubin** (μmol/L) | 3 (2-5.5) | 3 (1.5-3) | 4 (2-9) | 2 (1-7) |
| **AST** (IU/L) | 122 (94-137) | 94 (87-115) | 126 (108-154) | 138 (128-201) |
| **ALP** (IU/L) | 101 (63-134) | 75 (56-101) | 101 (58-116) | 134 (133-220) |
| **CK** (IU/L) | 368 (261-509) | 305 (272-479) | 503 (378-601) | 357 (241-368) |

**Supplementary Table E3.**
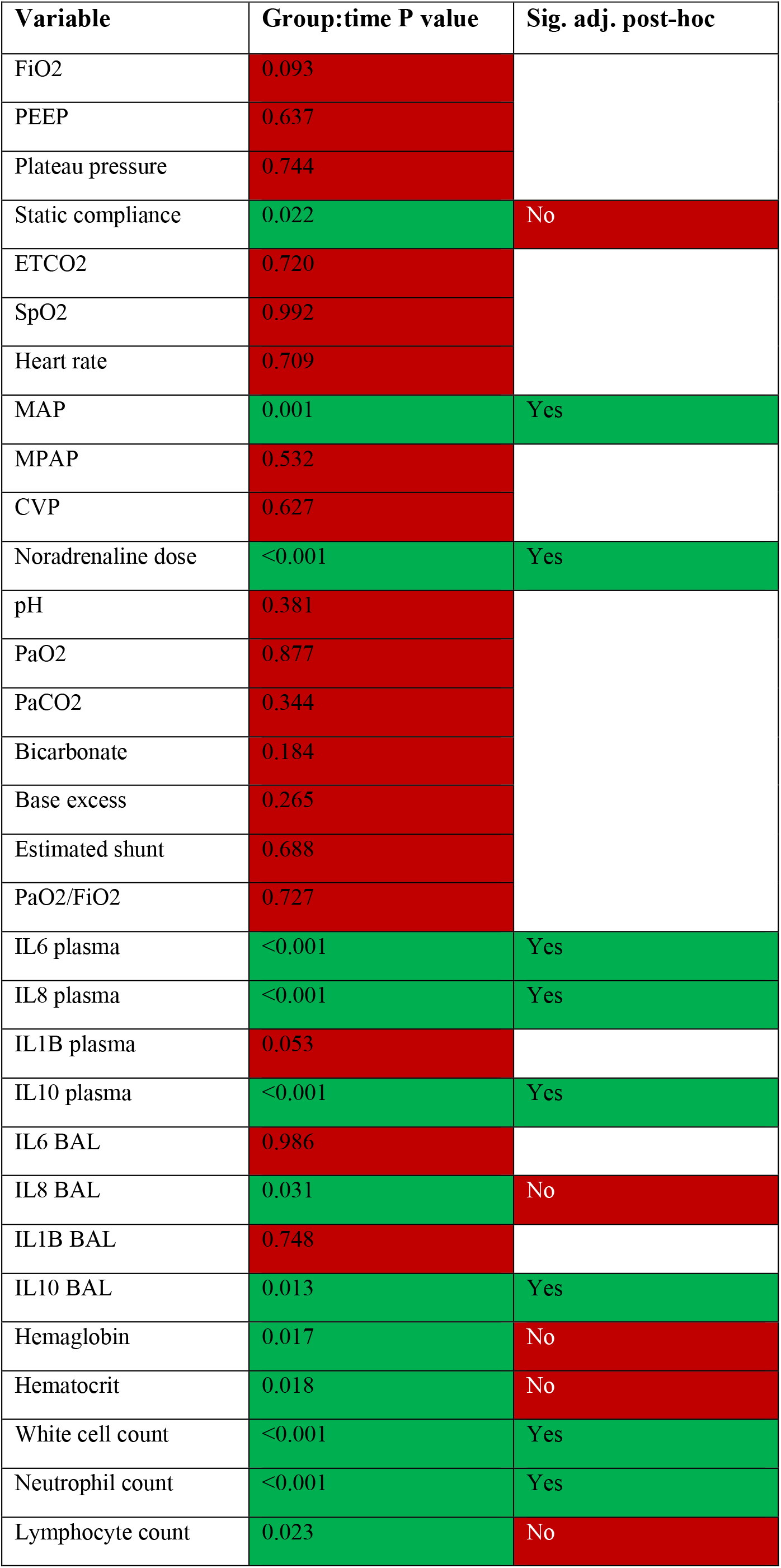
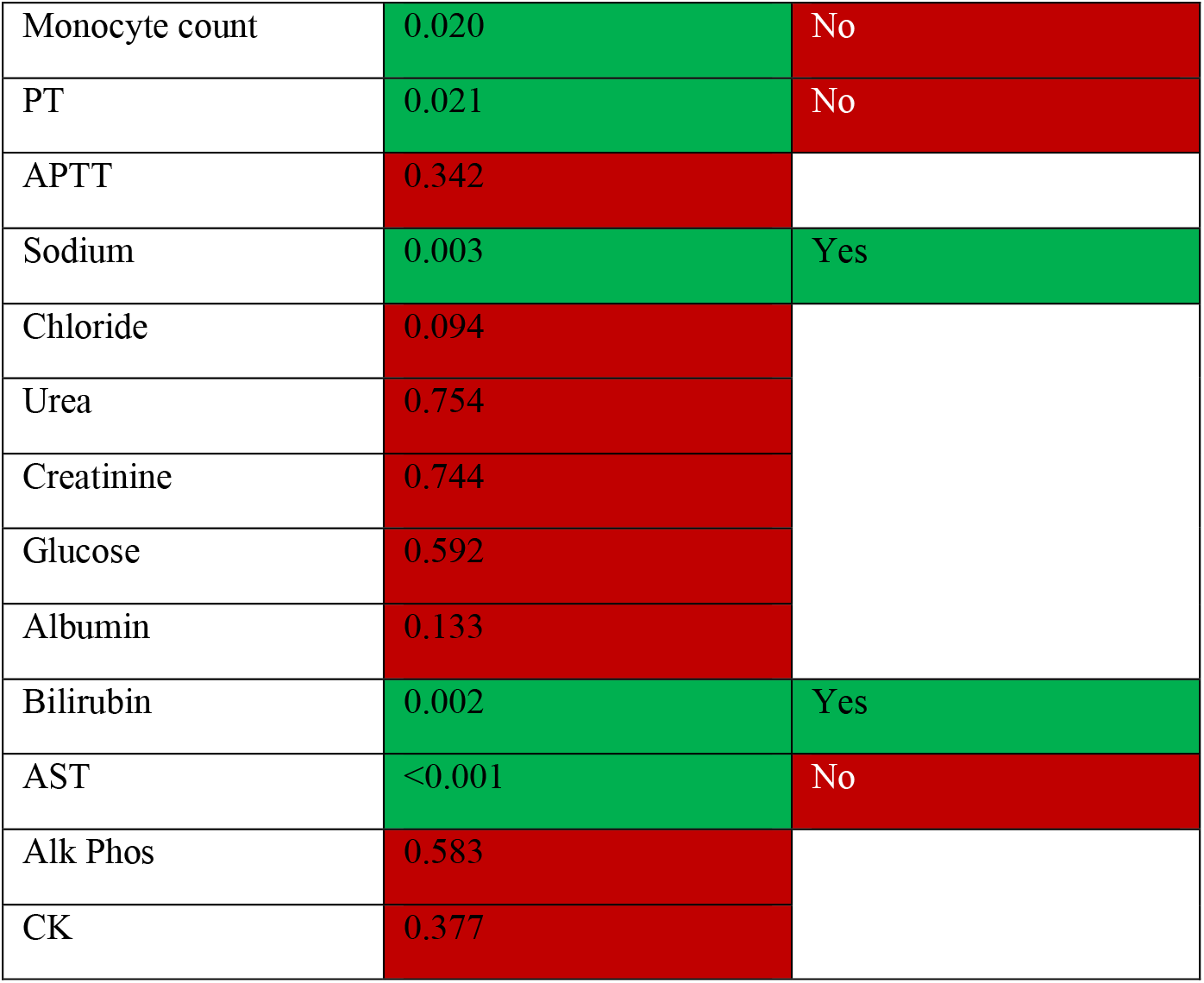
Linear mixed model group:time interactions.

| Variable | Group:time P value | Sig. adj. post-hoc |
| --- | --- | --- |
| FiO2 | 0.093 |  |
| PEEP | 0.637 |  |
| Plateau pressure | 0.744 |  |
| Static compliance | 0.022 | No |
| ETCO2 | 0.720 |  |
| SpO2 | 0.992 |  |
| Heart rate | 0.709 |  |
| MAP | 0.001 | Yes |
| MPAP | 0.532 |  |
| CVP | 0.627 |  |
| Noradrenaline dose | <0.001 | Yes |
| pH | 0.381 |  |
| PaO2 | 0.877 |  |
| PaCO2 | 0.344 |  |
| Bicarbonate | 0.184 |  |
| Base excess | 0.265 |  |
| Estimated shunt | 0.688 |  |
| PaO2/FiO2 | 0.727 |  |
| IL6 plasma | <0.001 | Yes |
| IL8 plasma | <0.001 | Yes |
| IL1B plasma | 0.053 |  |
| IL10 plasma | <0.001 | Yes |
| IL6 BAL | 0.986 |  |
| IL8 BAL | 0.031 | No |
| IL1B BAL | 0.748 |  |
| IL10 BAL | 0.013 | Yes |
| Hemaglobin | 0.017 | No |
| Hematocrit | 0.018 | No |
| White cell count | <0.001 | Yes |
| Neutrophil count | <0.001 | Yes |
| Lymphocyte count | 0.023 | No |
| Monocyte count | 0.020 | No |
| PT | 0.021 | No |
| APTT | 0.342 |  |
| Sodium | 0.003 | Yes |
| Chloride | 0.094 |  |
| Urea | 0.754 |  |
| Creatinine | 0.744 |  |
| Glucose | 0.592 |  |
| Albumin | 0.133 |  |
| Bilirubin | 0.002 | Yes |
| AST | <0.001 | No |
| Alk Phos | 0.583 |  |
| CK | 0.377 |  |

**Supplementary Table E4.** Post-hoc comparisons. Tukey’s honest significant differences test. P values adjusted using false discovery set at 5%.

| Timepoint | Group 1 | Group 2 | Mean difference | 95% CIs | adj. P value |
| --- | --- | --- | --- | --- | --- |
| 0 hrs | IT | IV | 27 | 7 – 47 | 0.012 |
| 0 hrs | IT | OA | 19 | 0 – 38 | 0.046 |
| 2 hrs | IT | OA | 20 | 1 – 38 | 0.034 |
| 6 hrs | IV | OA | 21 | 1 – 42 | 0.040 |

Noradrenaline dose, mcg/kg/min
| Timepoint | Group 1 | Group 2 | Mean difference | 95% CIs | adj. P value |
| --- | --- | --- | --- | --- | --- |
| 0 hrs | IT | OA | 0.211 | 0.015 – 0.408 | 0.034 |
| 0 hrs | IV | OA | 0.270 | 0.054 – 0.485 | 0.014 |
| 2 hrs | IT | IV | 0.486 | 0.125 – 0.848 | 0.008 |
| 2 hrs | IV | OA | -0.456 | -0.818 – -0.095 | 0.013 |

Plasma IL-6, ng/mL
| Timepoint | Group 1 | Group 2 | Mean difference | 95% CIs | adj. P value |
| --- | --- | --- | --- | --- | --- |
| 0 hrs | IT | OA | -1708 | -3306 – -109 | 0.036 |
| 1 hrs | IT | IV | -11964 | -23051 – -877 | 0.034 |
| 6 hrs | IT | OA | -21952 | -63392 – -5062 | 0.021 |

Plasma IL-8, ng/mL
| Timepoint | Group 1 | Group 2 | Mean difference | 95% CIs | adj. P value |
| --- | --- | --- | --- | --- | --- |
| 0 hrs | IT | IV | -1072 | -2000 – -144 | 0.023 |
| 2 hrs | IT | IV | 7796 | 1192 – 14400 | 0.020 |
| 2 hrs | IV | OA | -8561 | -15165 – -1957 | 0.011 |
| 6 hrs | IV | OA | -1022 | -1733 – -270 | 0.008 |

Plasma IL-10, ng/mL
| Timepoint | Group 1 | Group 2 | Mean difference | 95% CIs | adj. P value |
| --- | --- | --- | --- | --- | --- |
| 1 hrs | IT | IV | 17717 | 6547 – 28887 | 0.002 |
| 1 hrs | IV | OA | -14715 | -25885 – -3545 | 0.010 |
| 2 hrs | IT | IV | 37367 | 22441 – 52294 | <0.001 |
| 2 hrs | IV | OA | -38473 | -53400 – -23547 | <0.001 |
| 4 hrs | IT | IV | 22499 | 12493 – 32505 | <0.001 |
| 4 hrs | IV | OA | -23171 | -33177 – -13165 | <0.001 |
| 6 hrs | IT | IV | 25626 | 20461 – 30792 | <0.001 |
| 6 hrs | IV | OA | -26170 | -31335 – 21004 | <0.001 |

BAL IL-10, ng/mL
| Timepoint | Group 1 | Group 2 | Mean difference | 95% CIs | adj. P value |
| --- | --- | --- | --- | --- | --- |
| 1 hrs | IT | IV | 1264 | 138 – 2390 | 0.027 |
| 1 hrs | IT | OA | 1053 | 25 – 2081 | 0.044 |
| 2 hrs | IT | IV | 2267 | 365 – 4168 | 0.019 |

White cell count, $\times 10^9/L$
| Timepoint | Group 1 | Group 2 | Mean difference | 95% CIs | adj. P value |
| --- | --- | --- | --- | --- | --- |
| 4 hrs | IV | OA | 1.24 | 0 – 2.5 | 0.049 |

Neutrophil count, $\times 10^9/L$
| Timepoint | Group 1 | Group 2 | Mean difference | 95% CIs | adj. P value |
| --- | --- | --- | --- | --- | --- |
| 4 hrs | IV | OA | 0.97 | 0 – 1.9 | 0.034 |

Sodium, mmol/L
| Timepoint | Group 1 | Group 2 | Mean difference | 95% CIs | adj. P value |
| --- | --- | --- | --- | --- | --- |
| 1 hrs | IV | OA | -3 | -6 – 0 | 0.024 |
| 6 hrs | IT | IV | 4 | 1 – 7 | 0.008 |
| 6 hrs | IV | OA | -4 | -7 – -1 | 0.005 |

Bilirubin, $\mu\text{mol/L}$
| Timepoint | Group 1 | Group 2 | Mean difference | 95% CIs | adj. P value |
| --- | --- | --- | --- | --- | --- |
| 4 hrs | IT | OA | -2 | -4 – 0 | 0.042 |

**Supplementary Table E5.**
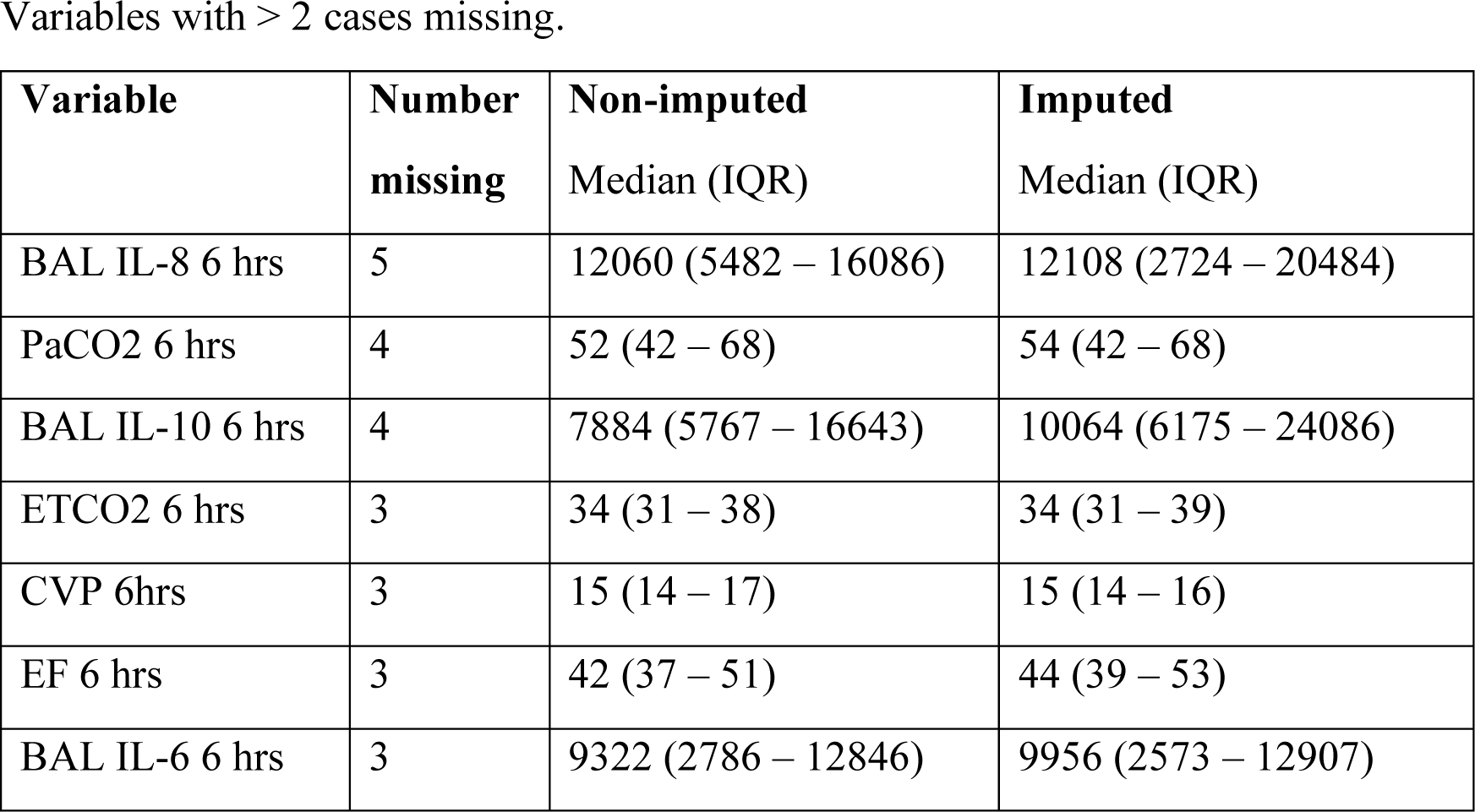
Data missingness.

Variables with > 2 cases missing.
| **Variable** | **Number**<br>**missing** | **Non-imputed**<br>**Median (IQR)** | **Imputed**<br>**Median (IQR)** |
| --- | --- | --- | --- |
| BAL IL-8 6 hrs | 5 | 12060 (5482 – 16086) | 12108 (2724 – 20484) |
| PaCO2 6 hrs | 4 | 52 (42 – 68) | 54 (42 – 68) |
| BAL IL-10 6 hrs | 4 | 7884 (5767 – 16643) | 10064 (6175 – 24086) |
| ETCO2 6 hrs | 3 | 34 (31 – 38) | 34 (31 – 39) |
| CVP 6hrs | 3 | 15 (14 – 17) | 15 (14 – 16) |
| EF 6 hrs | 3 | 42 (37 – 51) | 44 (39 – 53) |
| BAL IL-6 6 hrs | 3 | 9322 (2786 – 12846) | 9956 (2573 – 12907) |

**Supplementary Table E6.**
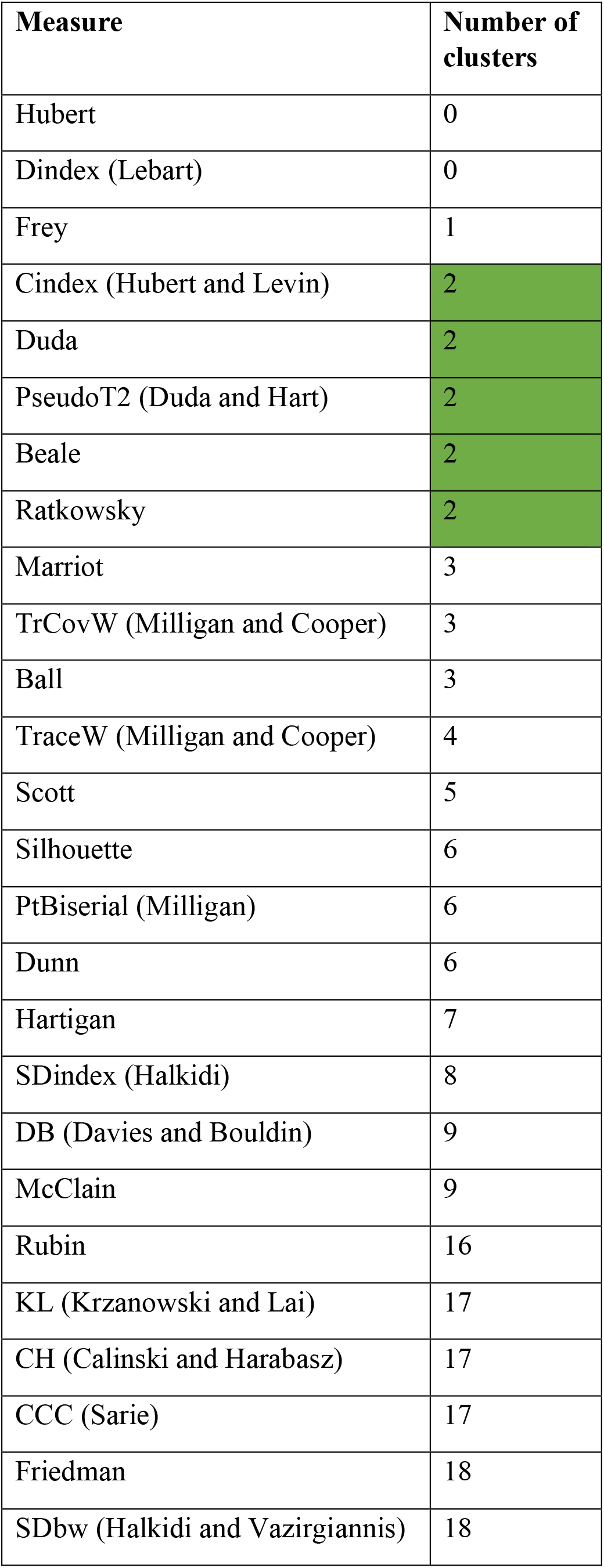
Measures of optimal cluster number

**Supplementary Figure E1.**
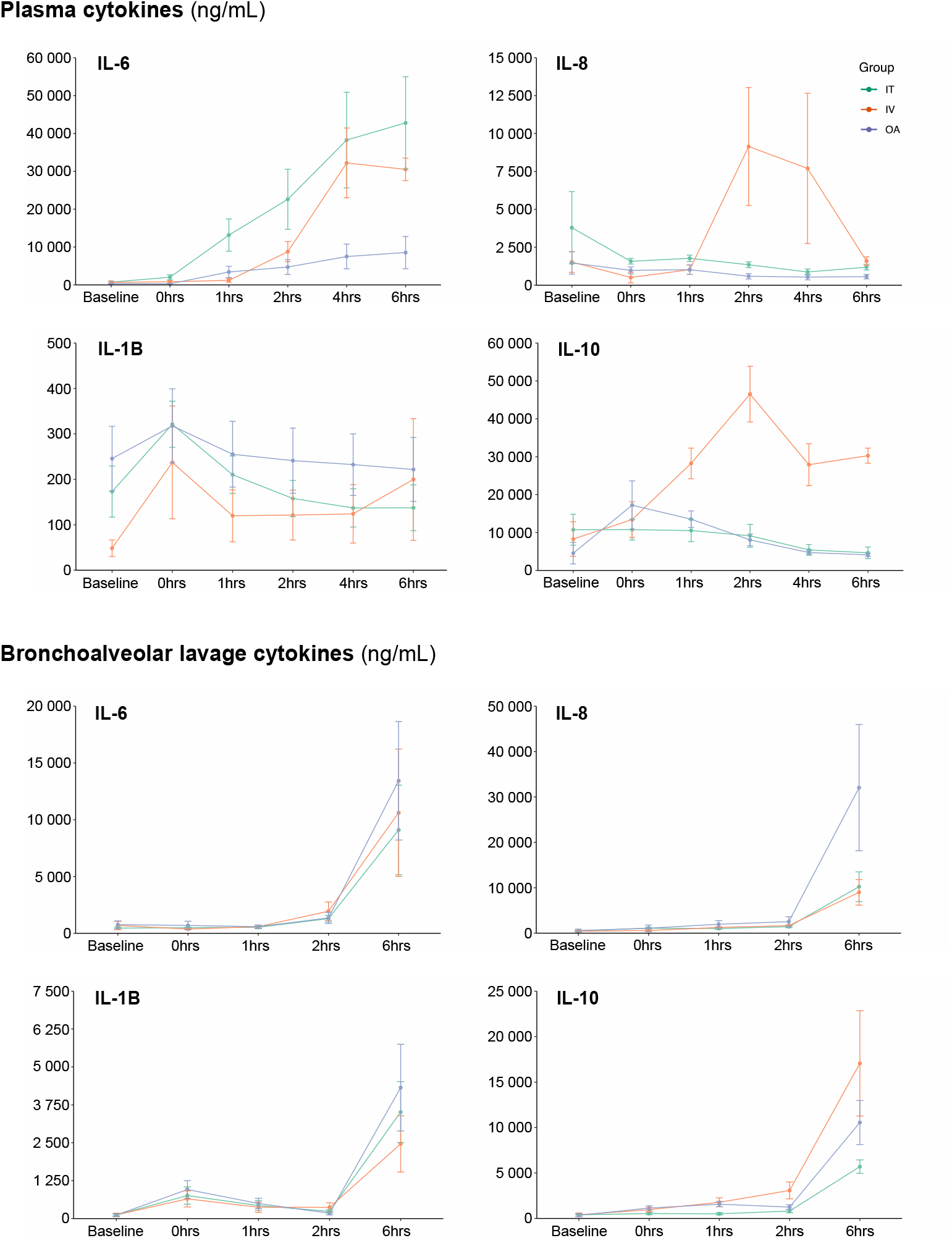
Plasma and bronchoalveolar lavage cytokine concentrations. Data are presented as mean and 95% confidence intervals.

**Supplementary Figure E2.**
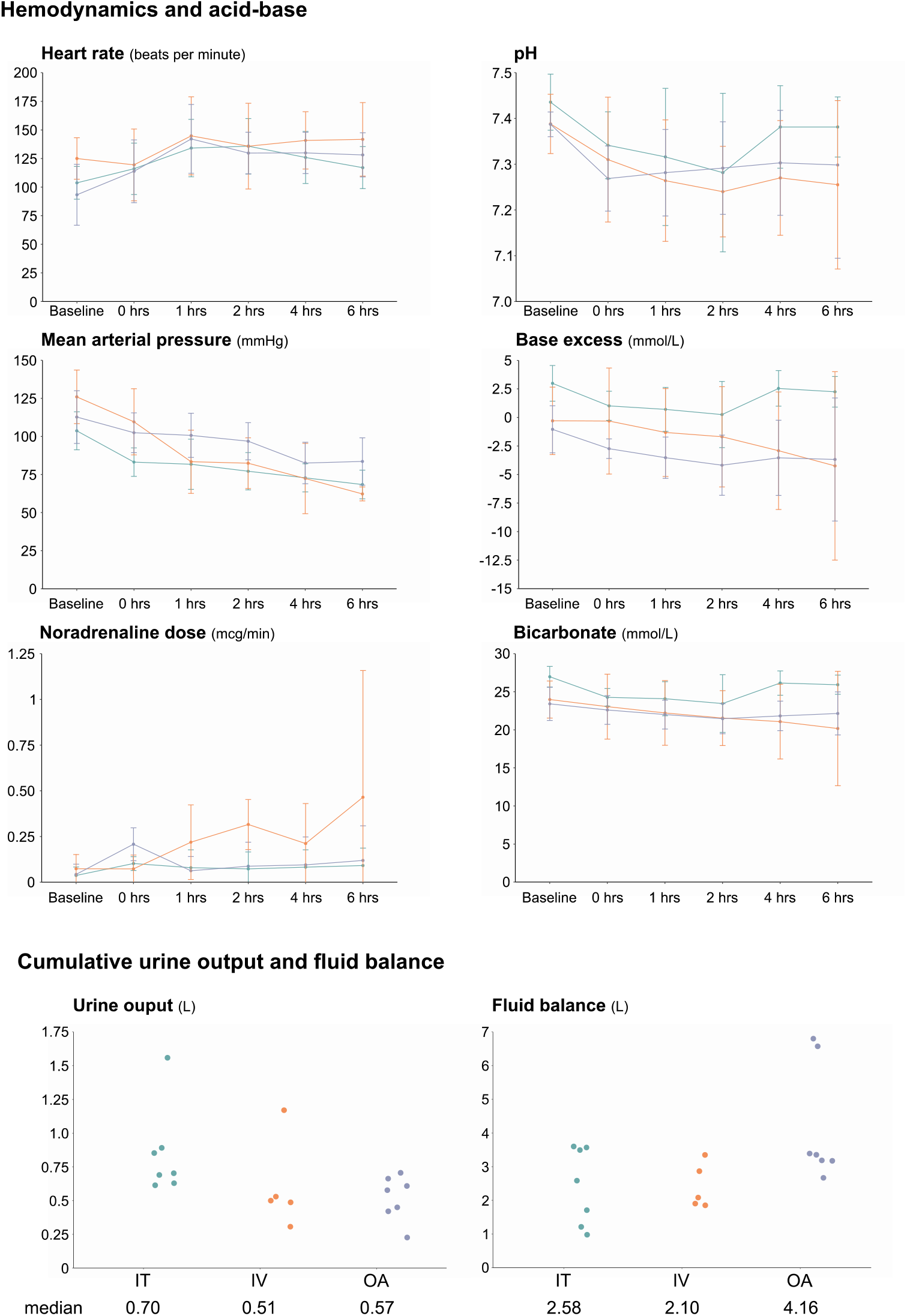
Hemodynamics, acid-base, and fluid balance. Data are presented as mean and 95% confidence intervals.

**Supplementary Figure E3.**
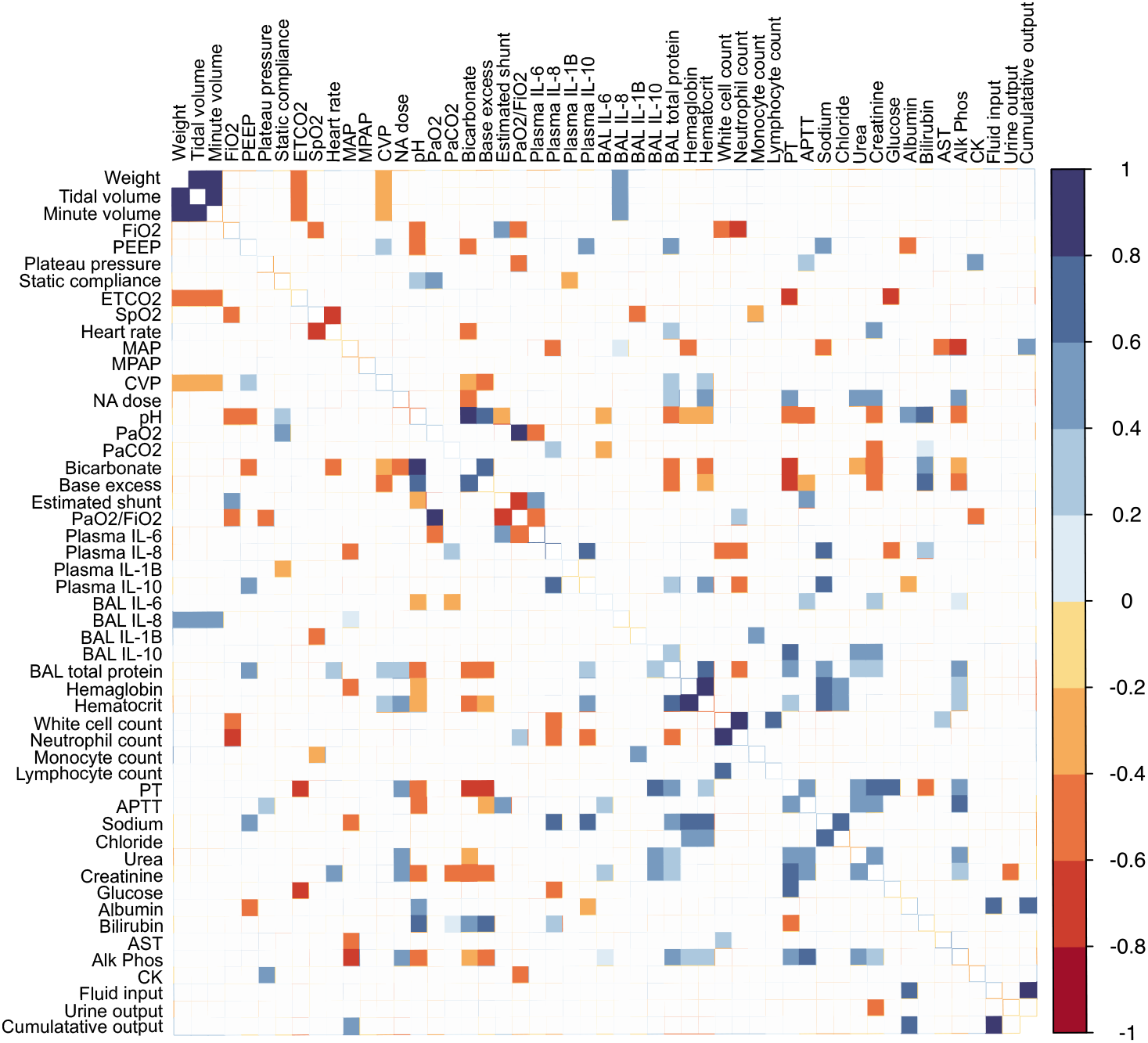
Correlation of variables. Spearman’s correlation. Pair-wise correlations with a p value > 0.05 are omitted.

**Supplementary Figure E4.**
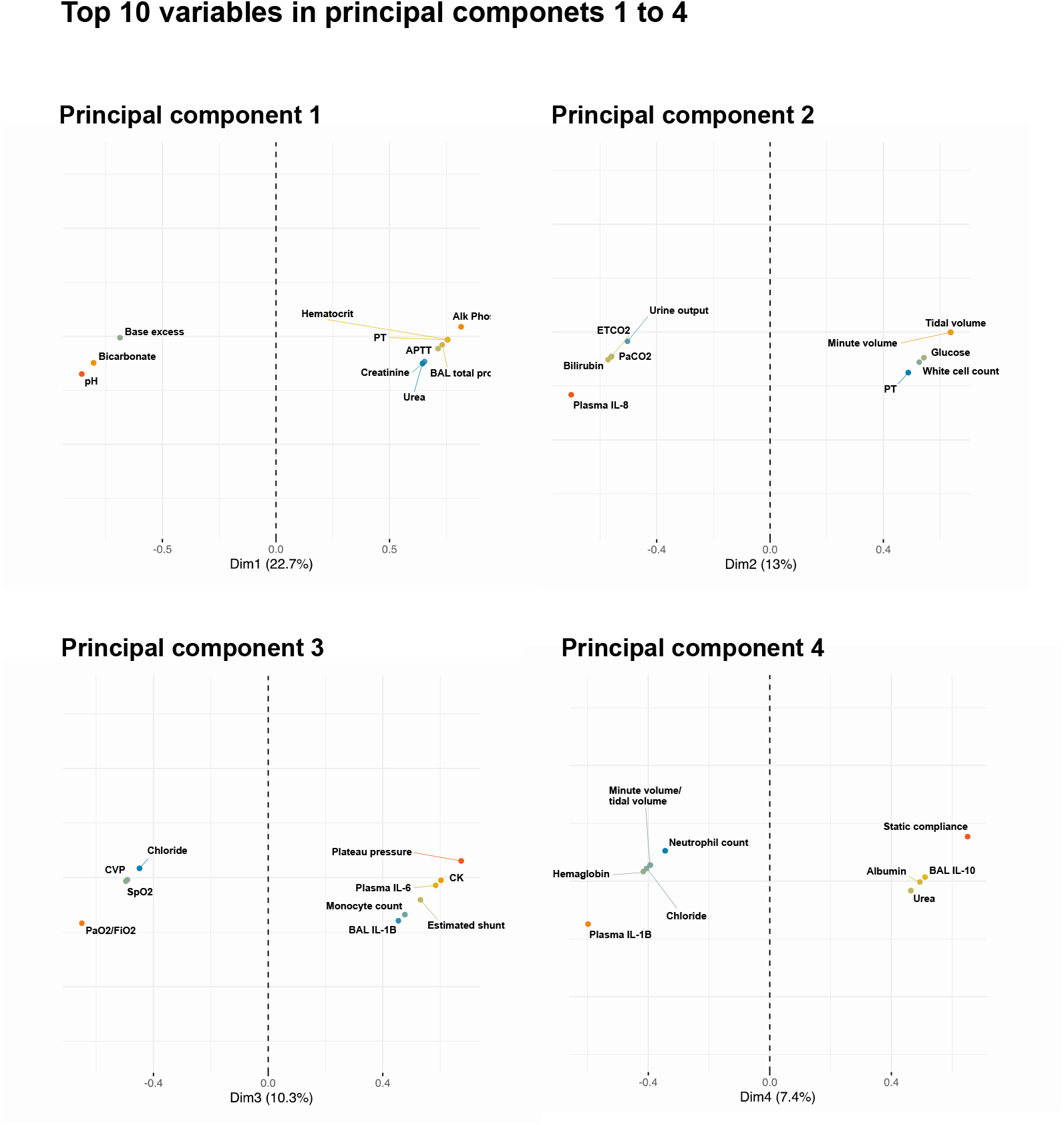
Contribution of variables to principal components 1-4. The ‘top 10’ variables describing each principal component are shown.

## References

1. Sinha P, Calfee CS. Phenotypes in acute respiratory distress syndrome: moving towards precision medicine. Curr Opin Crit Care 2019; 25: 12–20.

2. Calfee CS, Delucchi K, Parsons PE, Thompson BT, Ware LB, Matthay MA. Subphenotypes in acute respiratory distress syndrome: latent class analysis of data from two randomised controlled trials. Lancet Respir Med 2014; 2: 611–620.

3. Calfee CS, Delucchi KL, Sinha P, Matthay MA, Hackett J, Shankar-Hari M, McDowell C, Laffey JG, O’Kane CM, McAuley DF. Acute respiratory distress syndrome subphenotypes and differential response to simvastatin: secondary analysis of a randomised controlled trial. The Lancet Respir Med 2018; 6: 691–698.

4. Yehya N. Lessons learned in acute respiratory distress syndrome from the animal laboratory. Ann Transl Med 2019; 7: 503.

5. Seymour CW, Kerti SJ, Lewis AJ, Kennedy J, Brant E, Griepentrog JE, Zhang X, Angus DC, Chang C-CH, Rosengart MR. Murine sepsis phenotypes and differential treatment effects in a randomized trial of prompt antibiotics and fluids. Crit Care 2019; 23: 384–384.

6. National Health and Medical Research Council. Australian Code of Practice for the Care and Use of Animals for Scientific Purposes. 8th ed. Canberra: Australian Government; 2013.

7. Percie du Sert N, Hurst V, Ahluwalia A, Alam S, Avey MT, Baker M, Browne WJ, Clark A, Cuthill IC, Dirnagl U, Emerson M, Garner P, Holgate ST, Howells DW, Karp NA, Lidster K, MacCallum CJ, Macleod M, Petersen O, Rawle F, Reynolds P, Rooney K, Sena ES, Silberberg SD, Steckler T, Würbel H. The ARRIVE guidelines 2019: updated guidelines for reporting animal research. bioRxiv 2019: 703181.

8. Bouquet M, Passmore MR, See Hoe LE, Tung JP, Simonova G, Boon AC, Fraser JF. Development and validation of ELISAs for the quantitation of interleukin (IL)-1β, IL-6, IL-8 and IL-10 in ovine plasma. J Immunol Methods 2020; 486: 112835.

9. Husson F, Lê S, Pagès J. Exploratory multivariate analysis by example using R. Chapman and Hall/CRC; 2017.

10. Josse J, Husson F. Handling missing values in exploratory multivariate data analysis methods. Journal de la Société Française de Statistique 2012; 153: 79–99.

11. King EG, Nakane PK, Ashbaugh DG. The canine oleic acid model of fibrin localization in fat embolism. Surgery 1971; 69: 782–787.

12. Wang HM, Bodenstein M, Markstaller K. Overview of the pathology of three widely used animal models of acute lung injury. Eur Surg Res 2008; 40: 305–316.

13. Ballard-Croft C, Wang D, Sumpter LR, Zhou X, Zwischenberger JB. Large-animal models of acute respiratory distress syndrome. Ann Thorac Surg 2012; 93: 1331–1339.

14. Gonçalves-de-Albuquerque CF, Silva AR, Burth P, de Moraes IMM, Oliveira FMdJ, Younes-Ibrahim M, dos Santos MdCB, D’Ávila H, Bozza PT, Faria Neto HCdC, Faria MVdC. Oleic acid induces lung injury in mice through activation of the ERK pathway. Mediators Inflamm 2012; 2012: 956509–956509.

15. Moine P, McIntyre R, Schwartz MD, Kaneko D, Shenkar R, Le Tulzo Y, Moore EE, Abraham E. NF-kappaB regulatory mechanisms in alveolar macrophages from patients with acute respiratory distress syndrome. Shock 2000; 13: 85–91.

16. Gonçalves-de-Albuquerque CF, Silva AR, Burth P, Castro-Faria MV, Castro-Faria-Neto HC. Acute Respiratory Distress Syndrome: Role of Oleic Acid-Triggered Lung Injury and Inflammation. Mediators Inflamm 2015; 2015: 260465–260465.

17. Chen H, Bai C, Wang X. The value of the lipopolysaccharide-induced acute lung injury model in respiratory medicine. Expert Rev Respir Med 2010; 4: 773–783.

18. Wiener-Kronish JP, Albertine KH, Matthay MA. Differential responses of the endothelial and epithelial barriers of the lung in sheep to Escherichia coli endotoxin. J Clin Invest 1991; 88: 864–875.

19. Bos LD, Schouten LR, van Vught LA, Wiewel MA, Ong DSY, Cremer O, Artigas A, Martin-Loeches I, Hoogendijk AJ, van der Poll T, Horn J, Juffermans N, Calfee CS, Schultz MJ. Identification and validation of distinct biological phenotypes in patients with acute respiratory distress syndrome by cluster analysis. Thorax 2017; 72: 876–883.

20. Famous KR, Delucchi K, Ware LB, Kangelaris KN, Liu KD, Thompson BT, Calfee CS. Acute Respiratory Distress Syndrome Subphenotypes Respond Differently to Randomized Fluid Management Strategy. Am J Respir Crit Care Med 2017; 195: 331–338.

21. Sinha P, Delucchi KL, Thompson BT, McAuley DF, Matthay MA, Calfee CS. Latent class analysis of ARDS subphenotypes: a secondary analysis of the statins for acutely injured lungs from sepsis (SAILS) study. Intensive Care Med 2018; 44: 1859–1869.

22. Seymour CW, Kennedy JN, Wang S, Chang C-CH, Elliott CF, Xu Z, Berry S, Clermont G, Cooper G, Gomez H, Huang DT, Kellum JA, Mi Q, Opal SM, Talisa V, van der Poll T, Visweswaran S, Vodovotz Y, Weiss JC, Yealy DM, Yende S, Angus DC. Derivation, Validation, and Potential Treatment Implications of Novel Clinical Phenotypes for Sepsis. JAMA 2019; 321: 2003–2017.

23. Neyton LPA, Zheng X, Skouras C, Doeschl-Wilson A, Gutmann MU, Yau C, Uings I, Rao FV, Nicolas A, Marshall C, Wilson L-M, Baillie JK, Mole DJ. Molecular patterns in acute pancreatitis reflect generalizable endotypes of the host response to systemic injury in humans. Ann Surg 2020; published ahead of print doi: 10.1097/SLA.0000000000003974.

24. Vranas KC, Jopling JK, Sweeney TE, Ramsey MC, Milstein AS, Slatore CG, Escobar GJ, Liu VX. Identifying Distinct Subgroups of ICU Patients: A Machine Learning Approach. Crit Care Med 2017; 45: 1607–1615.

25. Jolliffe IT, Cadima J. Principal component analysis: a review and recent developments. Philos Trans Royal Soc A 2016; 374: 20150202.

26. Hümmeke-Oppers F, Hemelaar P, Pickkers P. Innovative Drugs to Target Renal Inflammation in Sepsis: Alkaline Phosphatase. Front Pharmacol 2019; 10: 919–919.

27. Baek SD, Kang J-Y, Yu H, Shin S, Park H-S, Kim M-S, Lee EK, Kim SM, Chang JW. Change in alkaline phosphatase activity associated with intensive care unit and hospital length of stay in patients with septic acute kidney injury on continuous renal replacement therapy. BMC Nephrol 2018; 19: 243–243.

28. Chang EM, Bretherick A, Drummond GB, Baillie JK. Predictive validity of a novel non-invasive estimation of effective shunt fraction in critically ill patients. Intensive Care Med Exp 2019; 7: 49–49.

29. Steinley D. K-means clustering: A half-century synthesis. Br J Math Stat Psychol 2006; 59: 1–34.

30. Castela Forte J, Perner A, van der Horst ICC. The use of clustering algorithms in critical care research to unravel patient heterogeneity. Intensive Care Med 2019; 45: 1025–1028.

## Supplement references

1. Checketts MR, Alladi R, Ferguson K, Gemmell L, Handy JM, Klein AA, Love NJ, Misra U, Morris C, Nathanson MH, Rodney GE, Verma R, Pandit JJ, Association of Anaesthetists of Great B, Ireland. Recommendations for standards of monitoring during anaesthesia and recovery 2015: Association of Anaesthetists of Great Britain and Ireland. Anaesthesia 2016; 71: 85–93.

2. Matute-Bello G, Downey G, Moore BB, Groshong SD, Matthay MA, Slutsky AS, Kuebler WM, Acute Lung Injury in Animals Study G. An official American Thoracic Society workshop report: features and measurements of experimental acute lung injury in animals. Am J Respir Cell Mol Biol 2011; 44: 725–738.

